# Noncanonical CDK4 signaling rescues diabetes in a mouse model by promoting beta cell differentiation

**DOI:** 10.1101/2022.10.19.512511

**Authors:** Rachel E. Stamateris, Rohit B. Sharma, Huguet Landa-Galván, Christine Darko, David Redmond, Sushil G. Rane, Laura C. Alonso

## Abstract

Expanding beta cell mass is a critical goal in the fight against diabetes. CDK4, an extensively characterized cell cycle activator, is required to establish and maintain beta cell number. Beta cell failure in the IRS2-deletion mouse type 2 diabetes model is in part due to loss of CDK4 regulator Cyclin D2. We set out to determine whether replacement of endogenous CDK4 with the inhibitor-resistant mutant CDK4-R24C rescued the loss of beta cell mass in Irs2-deficient mice. Surprisingly, not only beta cell mass but also beta cell dedifferentiation status was effectively rescued, despite no improvement in insulin sensitivity. *Ex vivo* studies in primary islet cells revealed a novel mechanism in which CDK4 intervened downstream in the insulin signaling pathway to prevent FOXO1-mediated transcriptional repression of critical beta cell transcription factor *Pdx1*. FOXO1 inhibition was not related to E2F1 activity, to FOXO1 phosphorylation, or even to FOXO1 subcellular localization, but rather was related to deacetylation of FOXO1 and reduced FOXO1 abundance. Taken together, these results demonstrate a novel differentiation-promoting activity of the classical cell cycle activator CDK4 and support the concept that beta cell mass can be expanded without compromising function.

## INTRODUCTION

An enduring goal to address the harms caused by diabetes is to increase endogenous pancreatic beta cell mass and function in people with insulin deficiency. Nutritional exposures such as hyperglycemia (1), high-fat diet (2, 3) and other nutrient excess paradigms (4) promote beta cell proliferation via mitogenic inputs that converge on downstream signaling to traverse the G1/S transition of the cell cycle (3, 5, 6). On the other hand, insulin secretory capacity can be lost through beta cell death (7) and dedifferentiation (8, 9). A critical barrier in the field is that strategies that increase replication may also lead to dedifferentiation (8–10).

Whole-body deletion of insulin receptor substrate (IRS)-2 in mice causes a type 2 diabetes (T2D)-like syndrome due to reduced beta cell function and mass in the face of marked insulin resistance (11, 12). Distal insulin signaling pathway member FOXO1 causes dedifferentiation in this model via suppression of key beta cell factor *Pdx1* (9, 13). IRS2-null beta cells also have an impaired proliferative response to glucose due to reduced induction of Cyclin D2, and restoring Cyclin D2 abundance rescues proliferation to normal levels (14). Cyclin D2, a known critical driver of postnatal beta cell expansion and beta cell compensation for insulin resistance (1, 15–19), binds to and activates cyclin-dependent kinase (CDK) family members CDK4/6 to hyperphosphorylate the Retinoblastoma (Rb) protein and derepress E2F transcription factors. CDK4 is critical for murine beta cells; deletion of CDK4 leads to a dramatic reduction in beta cell mass (20, 21). We postulated that if loss of CyclinD2/CDK4 activity in beta cells is a primary cause of diabetes in IRS2 null mice, then activating CDK4 in vivo (21) might counteract the diabetogenic phenotype in these mice. Specifically, we hypothesized that the *Cdk4-R24C* nucleotide substitution, which renders CDK4 uninhibitable by INK-family cell cycle inhibitors (22, 23), would rescue lost beta cell proliferation and mass in mice lacking IRS2.

Here we report that replacing both alleles of *Cdk4* with *Cdk4-R24C* rescued glucose intolerance in IRS2 null mice without improving insulin sensitivity. Beta cell mass and proliferation defects were rescued, as predicted. Surprisingly, *Cdk4-R24C* also corrected beta cell dedifferentiation, with full restoration of PDX1 and FOXO1 localization and cellular morphology. Intriguingly, CDK4 relieved FOXO1-mediated *Pdx1* repression, but the effect did not correlate with subcellular localization of FOXO1, nor did it require phosphorylation of FOXO1. Rather, the data were consistent with a model in which CDK4/CyclinD2 promoted histone deacetylase (HDAC) activity to suppress negative effects of FOXO1 in the setting of IRS2 deficiency. Taken together, this work highlights a novel role for the CDK4 kinase in the beta cell outside of cell cycle regulation, supporting the mature beta cell phenotype by derepressing *Pdx1* expression via the insulin signaling mediator FOXO1.

## RESULTS

### Homozygous replacement of *Cdk4* with *Cdk4-R24C* rescued diabetes in *Irs2*-null mice

To test whether activating CDK4 rescued lost beta cell mass in IRS2 null mice, we mated mice doubly heterozygous in all tissues (**Fig 1A**) for a loss-of-function allele of *Irs2* (12) and a gain-of-function allele of *Cdk4, Cdk4-R24C*, which contains a point mutation rendering CDK4 resistant to inhibition by INK-family inhibitors, knocked in at the endogenous *Cdk4* locus (21). Progeny of matings in which both dam and sire were *Irs2 +/-; Cdk4 wt/R24C* included the six genotypes of interest: *Irs2* wild-type (*+/+*) or knockout (*-/-*) that were wild-type (*wt/wt*), heterozygous (*wt/R24C*), or homozygous (*R24C/R24C*) for *Cdk4-R24C* at the *Cdk4* locus. We did not study mice heterozygous for *Irs2* deletion given their normal glucose homeostasis in our colony (14). Both males and females were analyzed.

**Figure 1.**
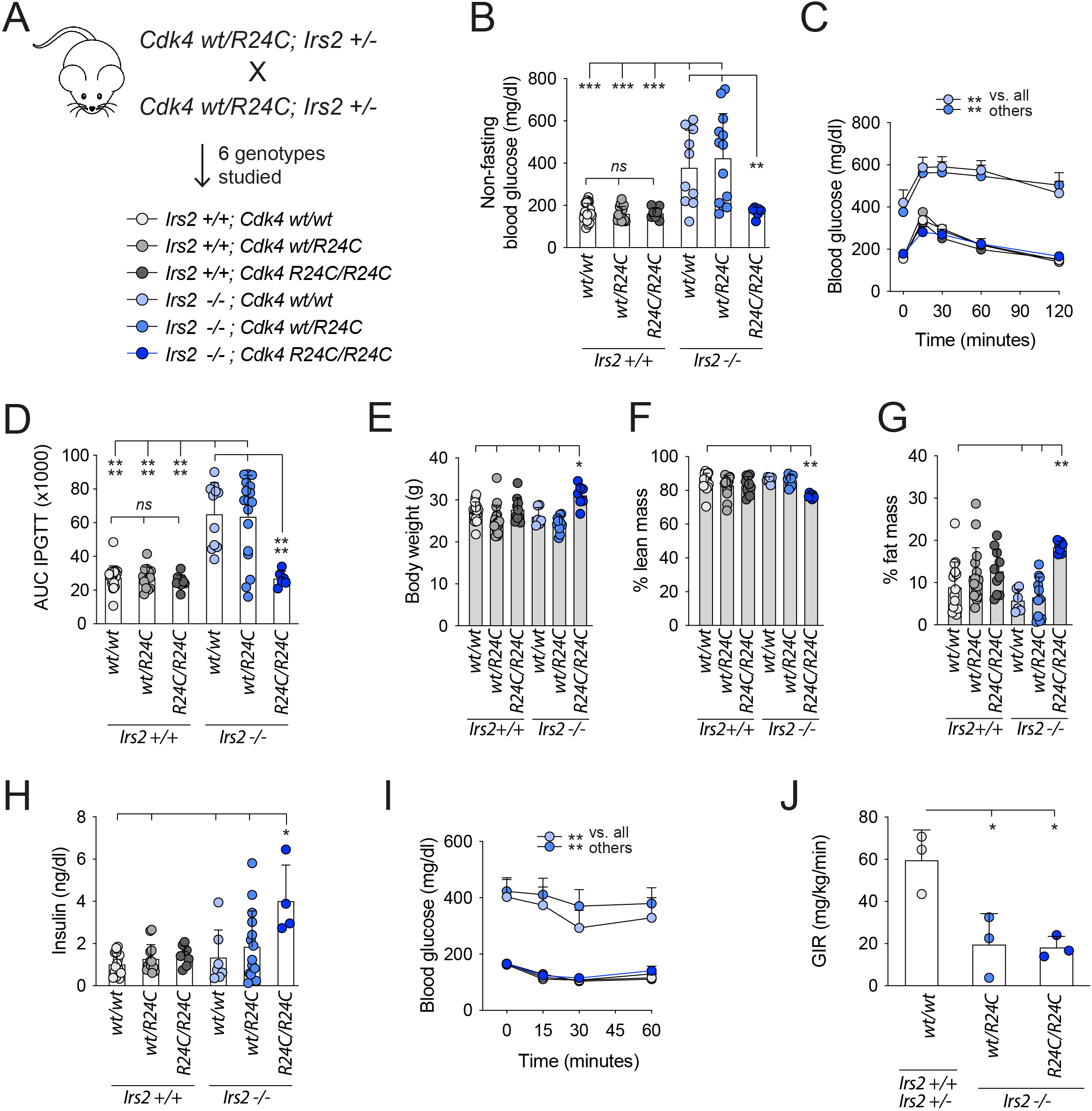
Homozygous replacement of *Cdk4* with *Cdk4-R24C* rescued diabetes in *Irs2*-null male mice. **A**: Breeding dams and sires doubly heterozygous for *Cdk4-R24C* (whole body knock-in) and the *Irs2-*null allele (whole-body knock-out) produced littermate 8-12 week experimental mice of the genotypes shown. *Irs2 +/-* progeny were not studied. **B-D:** non-fasting blood glucose (**B**), blood glucose time course (**C**) and AUC (**D**) after intraperitoneal glucose challenge. **E-G:** Body composition analysis by 1H-MRS Echo- MRI with body weight (**E**), % lean mass (**F**), and % fat mass (**G**). Circulating plasma insulin levels (**H**) were measured by ELISA. Glycemic response to intraperitoneal insulin challenge (**I**) was difficult to interpret due to markedly different baseline values. (**J**) Hyperinsulinemic euglycemic clamp showed that insulin resistance in *Irs2 -/-* mice was not rescued by homozygous *Cdk4-R24C*. Only male mice shown here, for females see **Suppl. Fig 1**. Number of replicates is shown for each panel except **C** (see **D**) and **I**, for which n=6-21. Statistics by one-way (**B, D, E, F, G, H, J**) or two-way (**C, I**) ANOVA with Tukey post-test. *p<0.05; **p<0.01; ***p<0.001; ****p<0.0001.

As expected (11, 12), male *Irs2-/-; Cdk4 wt/wt* mice had elevated nonfasting blood glucose (**Fig 1B**) and were markedly glucose intolerant (**Fig 1C-D**). Female *Irs2-/-; Cdk4 wt/wt* mice, as previously reported, had normal fasting and nonfasting blood glucose and normal glucose tolerance even when studied at a slightly older age or after 4 weeks of high fat feeding (**Suppl Fig 1A-G**). Remarkably, homozygous replacement of *Cdk4* with *Cdk4-R24C* in *Irs2* -/-males completely restored random non-fasting and post-challenge glucose curves to levels equivalent to nondiabetic controls (**Fig 1B-D**). *Cdk4* heterozygous (*wt/R24C*) replacement did not lead to discernible improvement in glucose tolerance in IRS2 null mice (**Fig 1B-D**). Thus, whole-body homozygous, but not heterozygous, replacement of *Cdk4* with *Cdk4-R24C* provided effective protection against diabetes in male mice lacking IRS2.

### *Cdk4-R24C* rescue of IRS2-null diabetes was not due to improved insulin sensitivity

In addition to islet growth benefits (20, 21, 24), CDK4 was reported to enhance insulin sensitivity in insulin responsive tissues such as adipose (25, 26). We hypothesized that restoration of glucose metabolism in *Irs2-/-* mice was due to improved insulin sensitivity, increased insulin secretory capacity, or both. *Irs2-/-;CDK4-R24C/R24C* males were heavier than littermate controls, with increased % fat mass and reduced lean mass (**Fig 1E-G**), an effect not observed in females (**Suppl Fig 1H-J**). Plasma insulin measurements confirmed hyperinsulinemia in *Irs2-/-;Cdk4-R24C/R24C* males (**Fig 1H**) suggesting rescue of insulin secretory capacity in the context of at least some residual insulin resistance. Insulin tolerance tests were difficult to interpret due to different baseline glucose levels (**Fig 1I**). Hyperinsulinemic euglycemic clamp revealed that the glucose infusion rate, low in *Irs2-/-* mice, was not rescued by homozygous *R24C* alleles, suggesting impaired insulin sensitivity in IRS2 null mice was not improved by *Cdk4-R24C* (**Fig 1J**). The increased fat mass in *Irs2-/-;Cdk4-R24C/R24C* mice (**Fig 1G**) could be consistent with enhanced insulin-responsive lipogenesis in adipocytes, but if present this was insufficient to rescue whole-body insulin resistance. Taken together, these data suggest that the rescue of diabetes in IRS2 null mice by *Cdk4-R24C* was due to correction of insulin deficiency rather than insulin responsiveness.

### *Cdk4-R24C* restored beta cell mass and increased beta cell proliferation in IRS2-null male mice

Since homozygous *Cdk4-R24C* alleles rescued glucose intolerance by correcting insulin deficiency, we predicted that beta cell function, mass, or both were restored in *Irs2-/-;Cdk4-R24C/R24C* mice. As expected, pancreas sections showed that islets in *Irs2-/-;Cdk4-wt/wt* mice had less immunoreactivity for insulin than controls. In contrast, *Irs2-/-;Cdk4-R24C/R24C* mice had fully rescued islet morphology and insulin staining (**Fig 2A**). Pancreas weight was similar among genotypes (**Fig 2B**), but beta cell mass estimated using a ratiometric approach showed a complete rescue of %insulin-stained area and beta cell mass in *Cdk4-R24C* homozygous, but not heterozygous, mice (**Fig 2C-D**). Since CDK4 is a positive regulator of the cell cycle, we hypothesized that the expanded beta cell mass in *Cdk4*-*R24C* homozygous mice was due to increased beta cell proliferation. In pancreas sections, the percent beta cells with nuclear BrdU (**Fig 2E-F**) or pHH3 (**Fig 2G-H**) labeling was increased in *Cdk4-R24C* homozygous mice over diabetic controls. Proliferation was only significantly increased by *Cdk4-R24C* in the IRS2-null context, suggesting the proliferation was amplified by insulin resistance. Thus, expanded beta cell mass due to increased proliferation likely contributed to the diabetes rescue.

**Figure 2.**
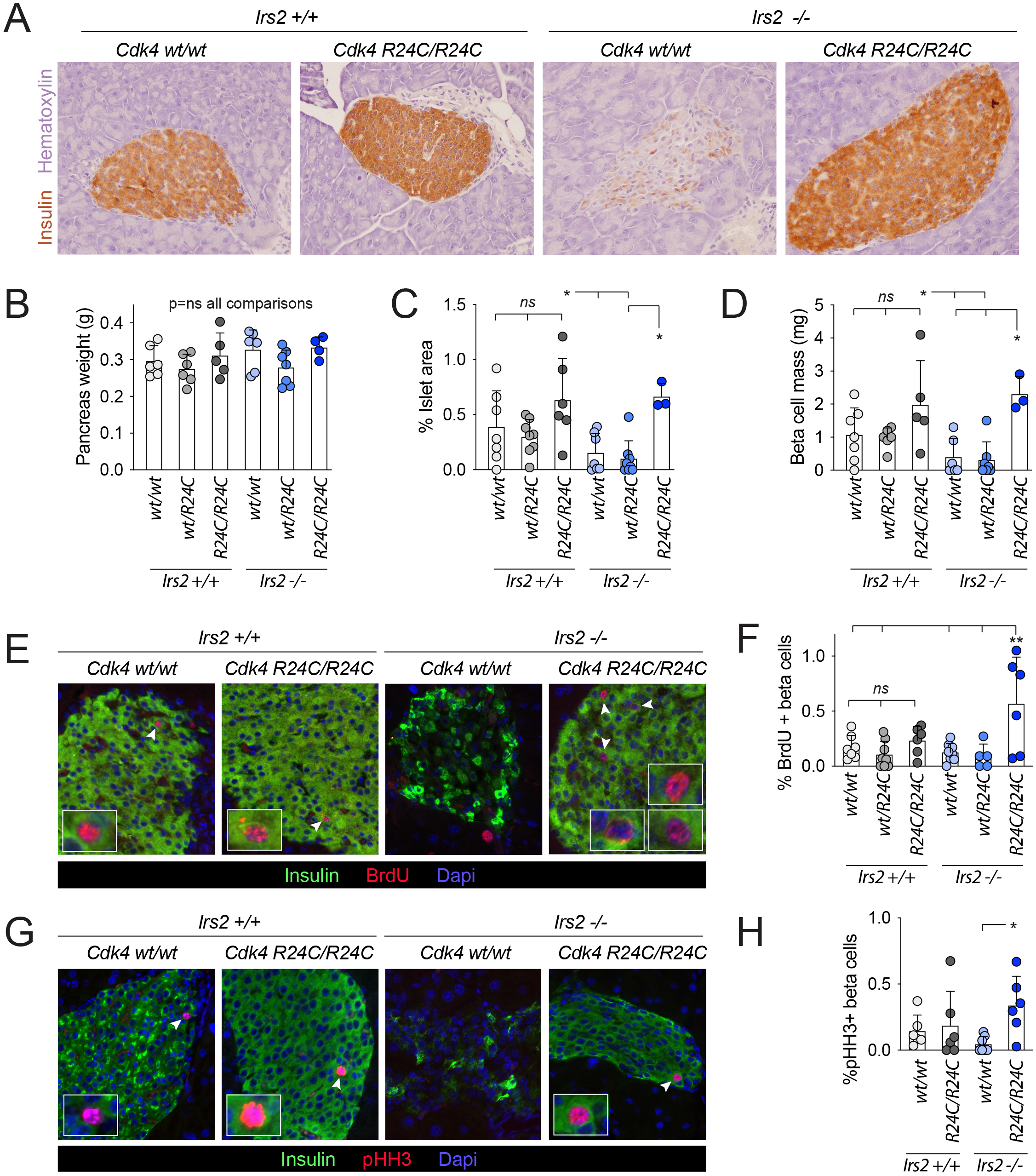
*Cdk4-R24C* rescued beta cell mass and proliferation in *Irs2*-null mice. **A**: Pancreas of adult male mice immunostained for insulin (brown) and counterstained with hematoxylin (purple). **B-D**: Wet weight of dissected pancreas prior to fixation (**B**) was multiplied by the % islet area (**C**) to estimate beta cell mass (**D**). **E-F**: Images from BrdU (red), insulin (green) and Dapi (blue) stained sections (**E**) were used to quantify the percent of insulin+ cells that had BrdU+ nuclei (**F**). **G-H**: Images from pHH3 (red), insulin (green) and Dapi (blue) stained sections (**G**) were used to quantify the percent of insulin+ cells that had pHH3+ nuclei (**H**). Number of replicates is shown for each panel. Statistics (**B, C, D, F, H**) by one-way ANOVA with Tukey post-test. *p<0.05; **p<0.01; ***p<0.001; ****p<0.0001.

### *Cdk4-R24C* restored islet morphological defects in IRS2 null mice

Although proliferation was statistically increased in *Irs2-/-;Cdk4-R24C/R24C* mice compared to *Irs2-/-* controls, the overall frequency of proliferation remained low and seemed insufficient to explain the marked improvement in islet morphology and insulin content. To better assess islet architecture, we performed immunofluorescence for insulin and glucagon (**Fig 3A**). *Irs2-/-;Cdk4 wt/wt* islets stained poorly for insulin, with reduced staining intensity in general and marked heterogeneity among beta cells compared to controls. Islet architecture was also disrupted in *Irs2* null islets, with alpha cells no longer restricted to the periphery but instead intermixed with beta cells (**Fig 3A**). Many of the insulin-low cells were also negative for glucagon, suggesting impaired beta cell insulin production rather than cell loss with resulting islet collapse, consistent with the previously demonstrated beta cell dedifferentiation reported in this model (13, 27). Remarkably, the quality and intensity of insulin staining, and the islet architecture, were completely restored to normal levels in mice bearing homozygous *Cdk4-R24C* alleles (**Fig 3A**).

**Figure 3.**
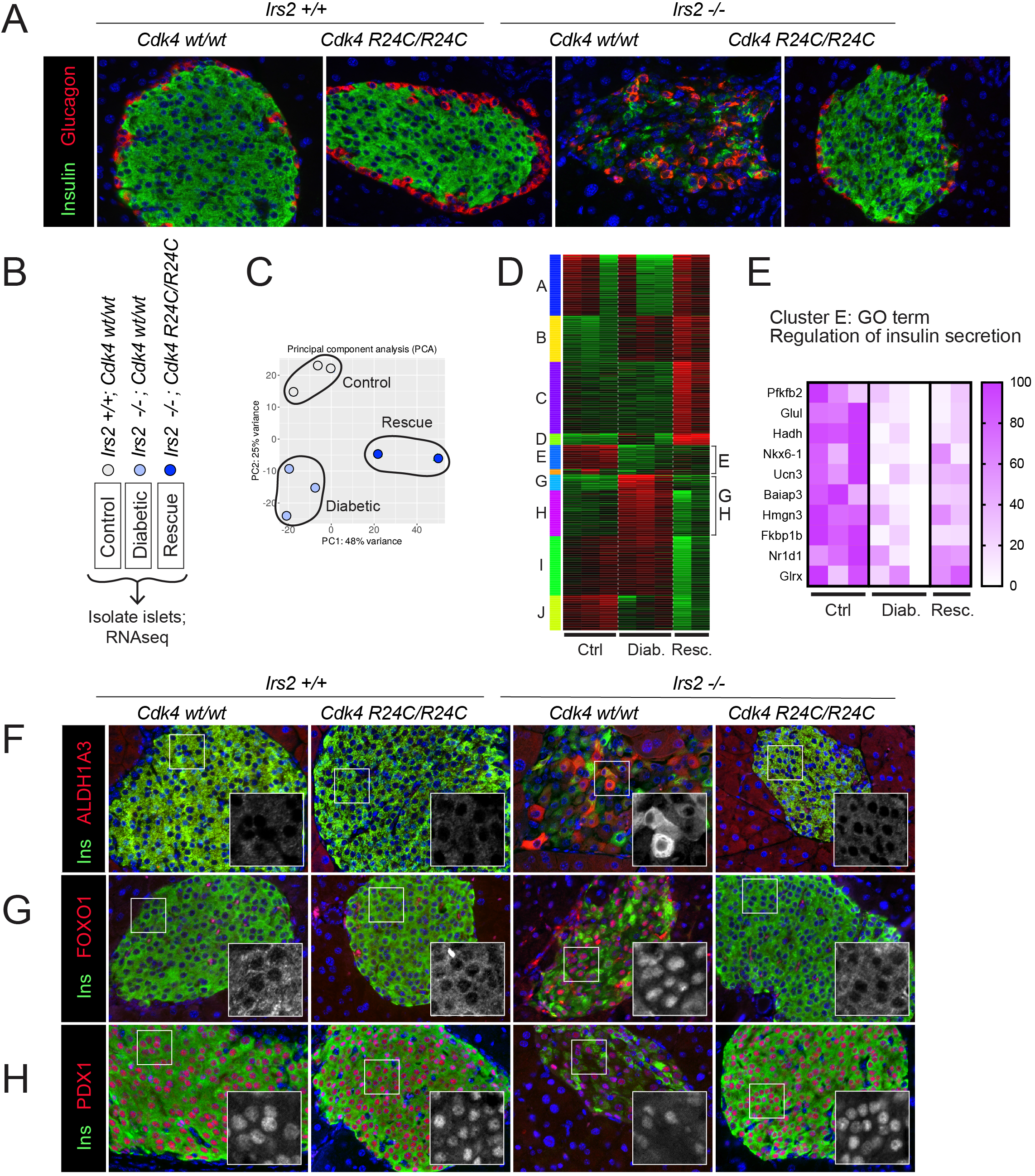
*Cdk4-R24C* corrected beta cell dedifferentiation in *Irs2-*null islets. **A**: Pancreas sections stained for insulin (green) and glucagon (red). **B-E**: Bulk RNA sequencing of islets isolated from Control (*Irs2+/+*;*Cdk4*-*wt/wt)*, Diabetic *(Irs2*-*/-*;*Cdk4*-*wt/wt)*, and Rescue (*Irs2*-*/-*;*Cdk4*-*R24C/R24C)* genotypes. Principal component analysis showed clean partitioning of the genotypes (**C**). K-means clustering (**D**) revealed three clusters in which mRNA abundance was disrupted during decompensation (Diabetic) and partially recovered in (Rescue). Cluster E, containing genes that were reduced in *Irs2*-*/-*;*Cdk4*-*wt/wt* (Diabetic) mice and partially restored in *Irs2*-*/-*;*Cdk4*-*R24C/R24C* (Rescue) mice, shown in the heat map in (**E**) mapped to a single GO term: Regulation of insulin secretion. **F-H**: Pancreas sections immunostained for insulin (green), dapi (blue) and ALDH1A3 (**F**), FOXO1 (**G**) or PDX1 (**H**) (red) confirm beta cell dedifferentiation signature in *Irs2-/-* male mice was rescued by homozygous *Cdk4-R24C*. Greyscale insets reflect red channel immunofluorescence in **F-H**.

### Gene signature analysis suggested rescue of beta cell function

To dissect the mechanism by which *R24C-R24C* rescued IRS2-null islet function, we used RNA sequencing to assess gene expression level changes. Whole islets were isolated from adult mice of control (*Irs2+/+*;*Cdk4*-*wt/wt)*, diabetic *(Irs2*-*/-*;*Cdk4*-*wt/wt)*, and rescue (*Irs2*-*/-*;*Cdk4*-*R24C/R24C)* genotypes (**Fig 3B**) and sent for library generation and sequencing. The samples cleanly partitioned based on principal component analysis (**Fig 3C**). Intriguingly, the major GO terms determining sample partitioning in principal components 1 and 2 involved RNA processing and peptide translation (**Suppl. Fig 2A**). Differential expression analysis revealed a number of differences between diabetic islets and controls (566 genes up, 327 genes down) and between rescue islets and diabetic islets (1064 up, 1115 down) (**Suppl. Fig 2B-D**). K-means clustering identified genes for which expression was reduced in diabetic islets and regained in rescue, or expression was increased in diabetic islets and restored to normal levels in rescue. 3 clusters (E, G, H) matched these patterns (**Fig 3D**). GO term mapping of genes increased in diabetic islets and corrected in rescue islets (clusters G-H) included protein glycosylation and response to ER stress. Cluster E, which contains genes lost in diabetic islets and partially regained in *R24C/R24C* rescued islets, mapped to a single GO term: Regulation of insulin secretion (**Fig 3E**). These data, though weakened by the caveats of bulk sequencing, inhomogeneous cell populations, and differences in the metabolic environment from which the samples were taken, supported the histological impression that *R24C/R24C* might improve beta cell function in IRS2-null mice (13, 27).

### *Cdk4-R24C* corrected beta cell dedifferentiation in IRS2 null mice

To assess dedifferentiation, we stained pancreas sections for aldehyde dehydrogenase 1 isoform A3 (ALDH1A3), a marker of failing beta cells (28). Intriguingly, the strong ALDH1A3 labeling observed in *Irs2*-/-;*Cdk4 wt/wt* beta cells was corrected in *Irs2-/-; Cdk4 R24C/R24C* mice (**Fig 3F**). FOXO1, a transcription factor suppressed by insulin signaling, is derepressed in IRS2 null mice and drives beta cell failure (13, 27). We hypothesized that *CDK4-R24C* might prevent FOXO1 activation in this model, mitigating the deleterious FOXO1 suppression of *Pdx1* expression. To investigate whether the FOXO1-PDX1 axis was restored in *Irs2*-/-; *Cdk4 R24C/R24C* islets, we first stained for FOXO1. Confirming prior observations (13), FOXO1 immunostaining was abnormally localized in beta cell nuclei in *Irs2-/-; Cdk4 wt/wt* islets (**Fig 3G**), consistent with reduced AKT-mediated phosphorylation of FOXO1 in the absence of IRS2. As previously published, *Irs2-/-; Cdk4 wt/wt* islets also had reduced nuclear staining for PDX1 (**Fig 3H**). Intriguingly, *Cdk4 R24C* homozygosity restored both FOXO1 and PDX1 staining to wild-type appearance, with FOXO1 restricted to the cytoplasm and PDX1 robustly nuclear (**Fig 3G-H**). This result suggests a novel role for activated CDK4 in promoting distal insulin signaling in the beta cell, specifically in protecting against FOXO1 mediated beta cell dedifferentiation in the setting of IRS2 deficiency.

### CDK4 rescued *Pdx1* and *Neurod1* expression in starve conditions ex vivo

We developed an ex vivo mouse islet cell starvation model to test CDK4 impact on FOXO1 activation. For these studies, CDK4 activity was increased by overexpression of wild type CDK4 (“CDK4”), Cyclin D2, or both. Short-term (16 hour) starvation of dispersed islet cells led to a heterogenous, but consistent, increase in nuclear FOXO1 (**Fig 4A**) and gene expression changes consistent with active nuclear FOXO1 such as increased *Cnr1* and decreased *Il6r* and *Gpd2* (**Fig 4B**)(29). Proliferation markers *PCNA* and *Ki67* were both decreased with starvation (**Fig 4C**). Starvation suppressed *Pdx1* expression, mimicking in vivo *Irs2* deletion; other beta cell differentiation markers decreased (*Mafa* and *NeuroD1*) or were not affected (*Nkx6*.*1* and *Ngn3*; **Fig 4D**).

**Figure 4.**
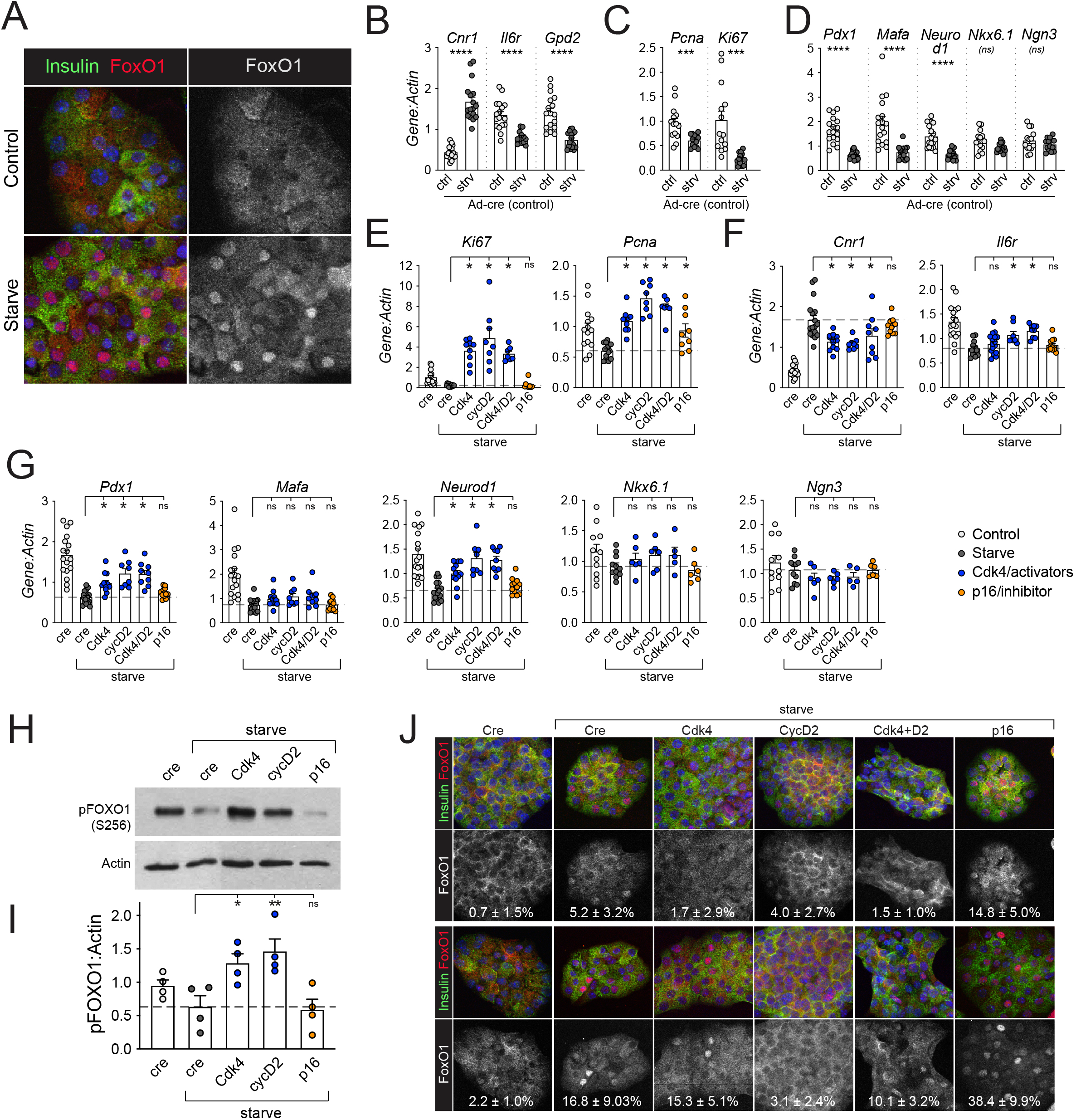
CDK4 suppresses starvation-induced FOXO1 activity in mouse islet cells. **A**: Dispersed mouse islet cells starved for 16 hours (2mM glucose, 0% FBS) were fixed, immunostained for FOXO1 (red), insulin (green) and Dapi (blue), and imaged by confocal microscopy. **B-G**: Dispersed mouse islet cells were transduced with the indicated adenoviruses, cultured in islet complete media for 72 hours with some wells starved for the last 16 hours, then lysed and analyzed by qPCR. With control (Ad-cre) virus, starvation altered known FOXO1 target gene expression in the expected directions (**B**), decreased proliferation markers (**C**) and reduced expression of some beta cell differentiation genes (**D**). **E-G:** Transduction with CDK4 activators (Ad-Cdk4, Ad-CyclinD2, Ad-Cdk4+Ad-CyclinD2), or CDK4 inhibitor (Ad- p16) showed that activating CDK4 rescued abundance of proliferation markers (**E**), rescued 2 of 3 FOXO1 targets (**F**), and rescued *Pdx1* and *Neurod1* expression but not *Mafa. Nkx6*.*1* and *Ngn3* were not changed by starvation or CDK4 overexpression (**G**). Data in **B-D** are the controls from **E-G**, presented separately for clarity. **H-I**: Ad-Cdk4 or Ad-Cyclin D2 increased phosphorylation of FOXO1 in dispersed mouse islet cells exposed to starve conditions (**H**), but Ad-p16 did not, quantified in (**I**). Confocal microscopy (**J**) with quantification of the percent beta cells with nuclear FOXO1 showed nuclear FOXO1 was only variably reduced by CDK4 activation. Top and bottom panels show two different biological replicates illustrating variability of nuclear FOXO1 abundance. Greyscale panels represent red channel (FOXO1) immunofluorescence. Dashed lines represent mean starve control condition. Number of replicates is shown for each panel. Statistics by T-test (**B-D**) or one-way ANOVA (**E-I**) with Tukey post-test. *p<0.05; **p<0.01; ***p<0.001; ****p<0.0001.

Using this starvation model, we tested whether CDK4 activation rescued the gene expression changes associated with nuclear FOXO1, by overexpressing CDK4 individually or in combination with Cyclin D2. As a negative control, we overexpressed p16, an INK family CDK4 inhibitor. As expected, the cell cycle activators increased proliferation markers *Ki67* and *Pcna*, while the inhibitor did not (**Fig 4E**). Interestingly, activating CDK4 partially rescued the starve-induced changes in two out of three FOXO1 targets (**Fig 4F**; *Gpd2* not rescued, not shown). Remarkably, overexpression of CDK4 with or without Cyclin D2 rescued *Pdx1* and *Neurod1* mRNA abundance (**Fig 4G**); *Mafa* levels were not rescued. On the other hand, the p16 CDK inhibitor did not rescue the FOXO1 target genes (**Fig 4F**), nor *Pdx1* or *Neurod1* (**Fig 4G**). *Nkx6*.*1* and *Ngn3*, which were not reduced by starvation (**Fig 4D**), were also not impacted by any of the cell cycle regulator combinations.

Since the immunofluorescence and mRNA data suggested that CDK4 inhibited FOXO1 activity, we tested whether CDK4 overexpression increased FOXO1 phosphorylation. By immunoblot, starvation decreased FOXO1 phosphorylation at S256, while overexpression of CDK4 or CyclinD2, but not p16, restored FOXO1 phosphorylation (**Fig 4H-I**). We hypothesized that CDK4 might directly phosphorylate FOXO1. Posttranslational modification prediction software (GPS 2.1) (30) predicted CDK4 consensus phosphorylation sites in FOXO1, some of which overlap with known AKT phosphorylation sites (data not shown). However, experiments using the AKT inhibitor MK-2206 showed that the increase in p-FOXO1 after CDK4 overexpression was dependent on AKT activation, suggesting an indirect mechanism. (**Suppl. Fig 3**).

### FOXO1 phosphorylation and activity did not correlate with subcellular localization

We hypothesized that CDK4-mediated alleviation of FOXO1 *Pdx1* suppression was related to the FOXO1 nuclear exclusion reported to follow phosphorylation (31, 32). We first tested whether increased FOXO1 phosphorylation resulted in FOXO1 cytoplasmic localization. However, in these experiments confocal microscopy showed that starvation (dephosphorylated FOXO1) caused only heterogeneously nuclear beta cell FOXO1 (**Fig 4J**). Further, overexpression of CDK4, which did increase FOXO1 phosphorylation (**Fig 4H-I**), only variably decreased the percentage of cells with visibly nuclear FOXO1 (**Fig 4J**). Intriguingly, p16 overexpression consistently increased the percent of beta cells with nuclear FOXO1 in this starvation model.

The variability in visibly nuclear FOXO1 from one experiment to the next, and the failure of CDK4 to consistently redistribute FOXO1 to the cytoplasm, suggested the possibility that CDK4 rescue of FOXO1-mediated *Pdx1* suppression might not require changing FOXO1 subcellular localization. To further explore the relationship between FOXO1 localization and expression of its transcription targets, we tested whether relocalizing FOXO1 to nuclei was sufficient to increase FOXO1 target gene expression. First, we treated mouse islet cells cultured in 15mM glucose with the nuclear export inhibitor Leptomycin B (33). Surprisingly, even in high glucose conditions where insulin signaling is fully activated in beta cells (14, 34), FOXO1 rapidly accumulated in the nucleus within 5-30 minutes of treatment with Leptomycin B, suggesting FOXO1 shuttles between the cytoplasm and nucleus even in nutrient replete conditions (**Fig 5A**). FOXO1 targets *Cnr1* and *Il6r* changed abundance in the expected direction; *Gpd2*, on the other hand, increased (**Fig 5B**). Despite robust nuclear accumulation of FOXO1, FOXO1 targets *Pdx1* and *Neurod1* did not decrease, but rather significantly increased (**Fig 5C**).

**Figure 5.**
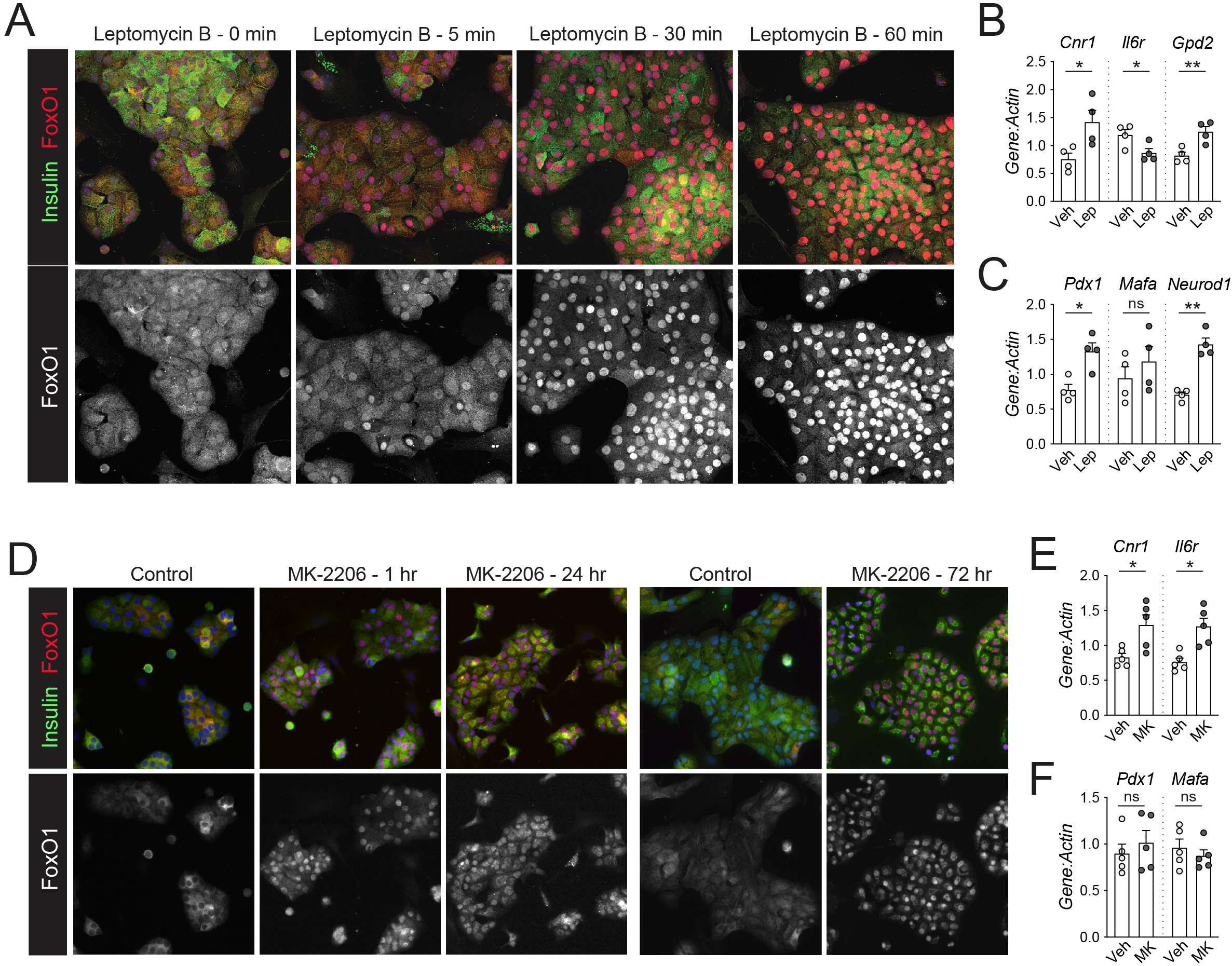
Forced nuclear accumulation of FOXO1 is not sufficient to repress *Pdx1*. **A**: Dispersed mouse islet cells cultured in ICM with 15mM glucose were treated with nuclear export inhibitor Leptomycin B (100nM) for the indicated duration, then fixed, immunostained for insulin (green), FOXO1 (red) and DAPI (blue), and imaged by confocal microscopy. The red (FOXO1) channel is displayed separately below. **B- C**: Dispersed mouse islet cells treated with Leptomycin B for 24 hours were analyzed by qPCR for FOXO1 targets (**B**) or beta cell maturation genes (**C**). **D-F**: Dispersed mouse islet cells cultured in ICM with 15mM glucose were treated with AKT inhibitor MK-2206 (5uM) for the indicated durations, then fixed, immunostained as above, and imaged on a standard fluorescence microscope (**D**) or lysed for qPCR (72h) (**E-F**). (**A**) and (**D**) are representative images; number of replicates is shown for all other panels. Statistics (**B, C, E, F**) are by unpaired T-test. *p<0.05, **p<0.01.

Leptomycin B traps many factors in the nucleus, so these effects may be independent of FOXO1. To more specifically test the relationship between nuclear FOXO1 and *Pdx1* suppression we treated dispersed islet cells with MK-2206, a pan-AKT inhibitor (35). We expected that inhibiting the kinase primarily responsible for phosphorylating FOXO1 (31) would lead to nuclear FOXO1 and suppression of *Pdx1*. Indeed, inhibiting AKT led to rapid and sustained FOXO1 nuclear accumulation (**Fig 5D**). However, despite nuclear localization, FOXO1 still failed to suppress *Pdx1* or elicit other expected gene expression changes (**Fig 5E-F**). With the caveat that both of these approaches alter cellular biology beyond FOXO1 modulation, these results suggested that nuclear localization of FOXO1 is not sufficient to suppress *Pdx1* in mouse beta cells.

### CDK4 rescue of FOXO1-induced *Pdx1* repression is independent of FOXO1 phosphorylation or subcellular localization

Lack of influence of CDK4 on FOXO1 localization led us to question whether CDK4 derepression of *Pdx1* involved FOXO1. To directly test whether CDK4 could counteract FOXO1-mediated *Pdx1* suppression we overexpressed FOXO1 in dispersed mouse islet cells cultured in high glucose and examined the impact of CDK4 co-expression (**Fig 6A**). Consistent with prior reports, FOXO1 overexpression suppressed *Pdx1* and *Il6r* mRNA (**Fig 6B-C**); different from prior reports, other mRNAs such as *MafA, NeuroD1* and *Cnr1* were also decreased (13, 29, 36). Coexpression of CDK4 and Cyclin D2 led to a marked increase in *Pdx1* mRNA (**Fig 6C**), confirming that CDK4 inhibited FOXO1-mediated *Pdx1* repression. Despite overexpression, FOXO1 did not visibly accumulate in beta cell nuclei (**Fig 6D**), reconfirming the lack of correlation between nuclear FOXO1 localization and gene expression changes in this system.

**Figure 6.**
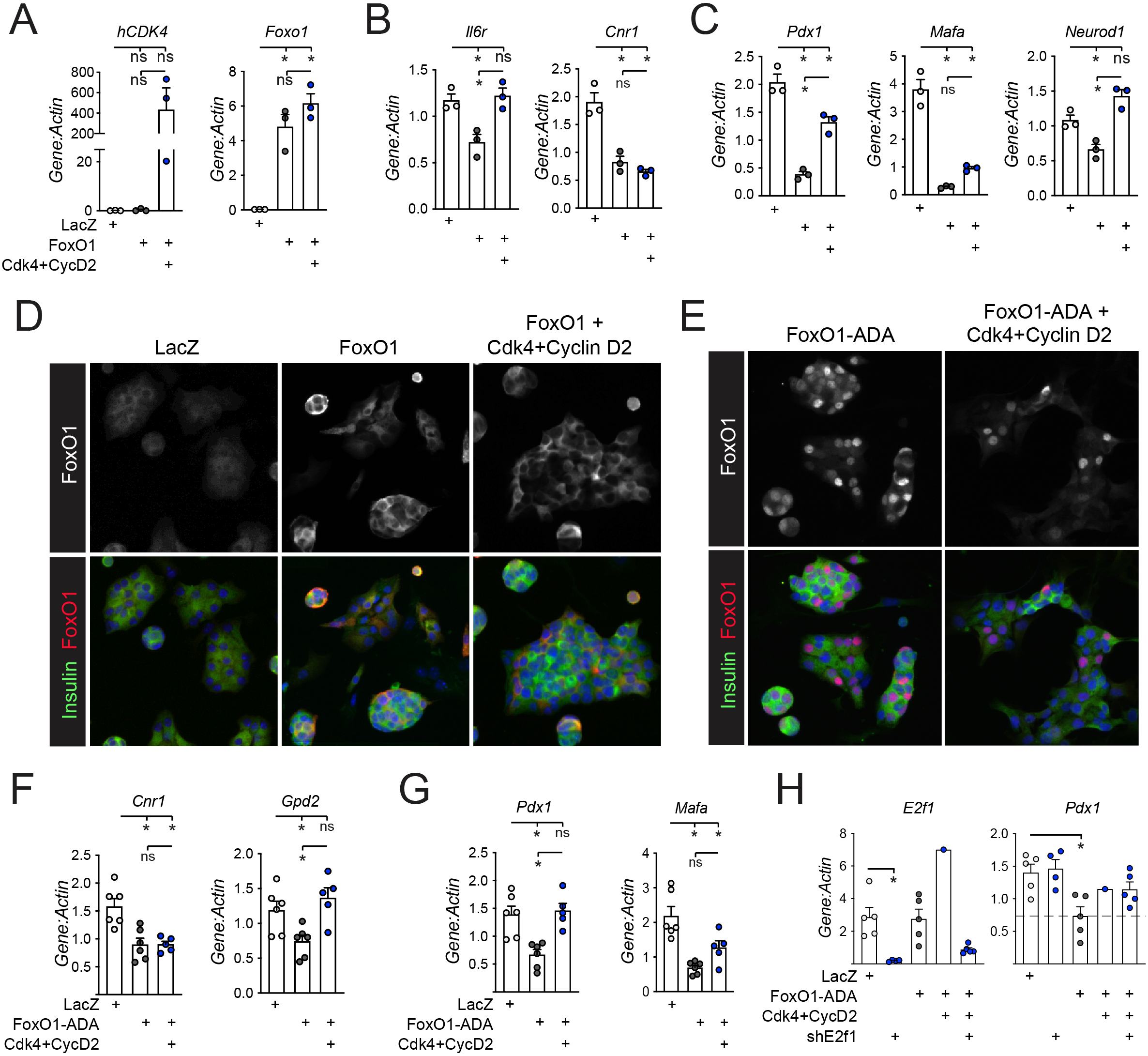
CDK4 rescues FOXO1-mediated *Pdx1* suppression even with an unphosphorylatable FOXO1-ADA mutant. **A-D**: Dispersed mouse islet cells were transduced with Ad-LacZ control, Ad-FOXO1, and/or Ad-CDK4 with Ad-Cyclin D2, cultured for 72 hours in ICM with 15mM glucose, then analyzed by qPCR for (**A**) *hCDK4* or *Foxo1* to assess transduction, (**B**) FOXO1 targets, or (**C**) beta cell differentiation genes; (**D**) parallel cultures on glass coverslips were immunostained for insulin (green), FOXO1 (red) and DAPI (blue) to assess FOXO1 localization in beta cells. **E-G**: Dispersed mouse islet cells were transduced with Ad-LacZ (control), Ad-FOXO1-ADA (unphosphorylatable mutant), and Ad-CDK4 + Ad-Cyclin D2, cultured for 72 hours in ICM with 15mM glucose, and subjected to (**E**) immunostaining for insulin (green), FOXO1 (red) and DAPI (blue) to assess FOXO1 localization in beta cells, or qPCR for (**F**) FOXO1 targets or (**G**) beta cell differentiation genes. **H**: Dispersed mouse islet cells transduced with the indicated adenoviruses for 72 hours in ICM with 15mM glucose were lysed for qPCR for *E2f1* or *Pdx1*. Number of replicates is shown for all qPCR panels. Statistics are by one-way ANOVA with Tukey post-test, *p<0.05.

To test whether phosphorylation of FOXO1 was required for CDK4-mediated derepression, we expressed the FOXO1-ADA mutant. FOXO1-ADA has serine to alanine/aspartic acid mutations in three AKT phosphorylation sites (T24A, S253D and S316A), making FOXO1 resistant to AKT phosphorylation and thus constitutively active and nuclear (37, 38). As expected, expressing FOXO1-ADA in dispersed mouse islet cells resulted in nuclear FOXO1 labeling (**Fig 6E**). FOXO1-ADA repressed FOXO1 targets *Cnr1* and *Gpd2* (**Fig 6F**) as well as *Pdx1* and *Mafa* (29) (**Fig 6G**). Surprisingly, co-expression of CDK4+CyclinD2 still restored *Pdx1* expression, suggesting that CDK4-mediated derepression of *Pdx1* does not require FOXO1 phosphorylation at AKT sites (**Fig 6G**). We explored whether CDK4 derepression of *Pdx1* expression involved canonical CDK activity via Rb phosphorylation and increased E2F1 activity, but found that knockdown of *E2f1* did not prevent CDK4 rescue of FOXO1-ADA mediated *Pdx1* suppression (**Fig 6H**). Taken together, these experiments suggest that although CDK4 overexpression increases FOXO1 phosphorylation in AKT-dependent fashion, the CDK4-mediated inhibition of FOXO1 repression of *Pdx1* and *Mafa* are independent of FOXO1 phosphorylation, nuclear localization, or E2F1 activation.

### CDK4 may modulate FOXO1 activity through deacetylation and promoting FOXO1 degradation

FOXO1 activity is regulated not only by phosphorylation but also by acetylation (36–41). Previous reports show CDKs can activate both histone acetyltransferases (HATs)(42) and histone deacetylases (HDACs)(43, 44). We tested whether modulating HAT or HDAC activity impacts FOXO1 suppression of *Pdx1* using small molecule inhibitors of HATs (MB-3 to inhibit GCN5 and C646 to inhibit p300/CBP) and HDACs (salermide to inhibit the sirtuins, sodium butyrate to inhibit class I/II HDACs, and vorinostat/SAHA as a general HDAC inhibitor)(**Fig 7A**). As previously demonstrated (45, 46), inhibiting p300 reduced *Ins1, Ins2* abundance (**Fig 7B**). Using the FOXO1-ADA mutant to eliminate influences on FOXO1 phosphorylation and localization, we found that inhibiting HATs tended to derepress *Pdx1* and *Mafa* to a similar degree as overexpression of CDK4/Cyclin D2 (**Fig 7C-D**). Conversely, inhibiting HDACs tended to have the opposite effect, impairing CDK4/Cyclin D2-induced derepression of *Pdx1* and *Mafa* (**Fig 7E-F**). These results suggest that CDK4/Cyclin D2 acts to either activate HDAC activity or repress HAT activity. Deacetylated FOXO1 has enhanced transcriptional activity (47, 48) but also increased degradation (36). Intriguingly, overexpression of CDK4/CyclinD2 increased phosphorylation of SIRT1, known to deacetylate FOXO1 (49–51)(**Fig 7G**). We attempted to assess FOXO1 acetylation status by immunoprecipitation but were unable to confidently quantify this parameter (data not shown). However, we did observe that CDK4/Cyclin D2 decreased total FOXO1 protein levels, consistent with enhanced degradation (**Fig 7H**). In sum, these results suggest that CDK4 promotes HDAC activity in islet cells, either of FOXO1 itself or other elements at the *Pdx1* promoter, to derepress FOXO1-induced *Pdx1* suppression and reduce FOXO1 abundance.

**Figure 7.**
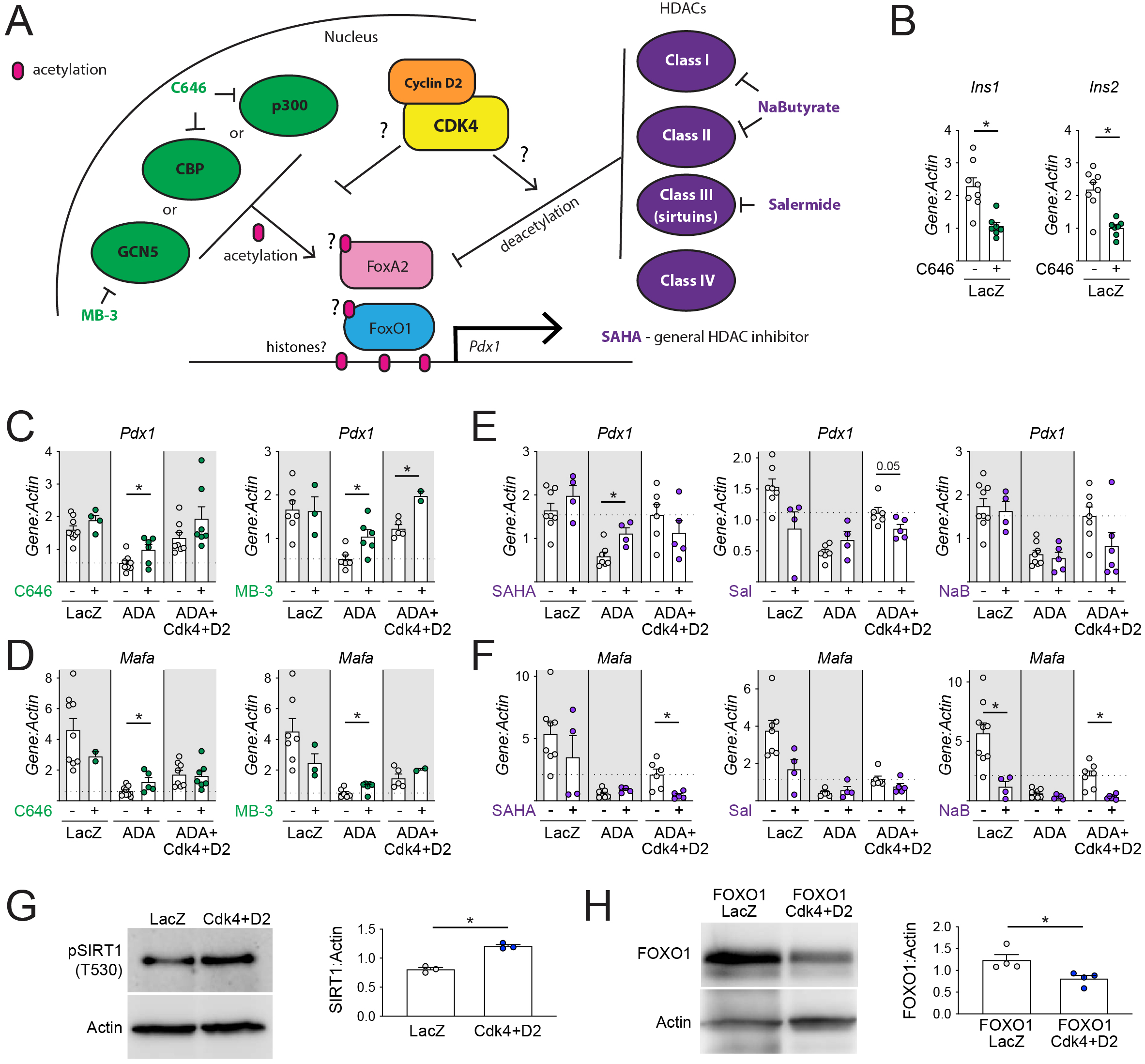
CDK4 rescue of FOXO1 *Pdx1* and *Mafa* suppression is modulated by acetylation. **A**: Model expressing the hypothesis that CDK4 influences acetylation of key players at the *Pdx1* locus, with diagram of histone acetyltransferases (HATs) and their inhibitors (green) and histone deacetylases (HDACs) and their inhibitors (purple). **C-F**: Dispersed mouse islet cells were transduced with the indicated adenoviruses, then cultured for 72 hours with or without the indicated HAT (**C-D**) or HDAC (**E-F**) inhibitors, then analyzed by qPCR for *Pdx1* or *Mafa*. **G-H**: Dispersed mouse islet cells were transduced with the indicated adenoviruses, cultured for 72 hours in ICM with 15mM glucose, then lysed and analyzed by immunoblot. Number of replicates is shown for all panels. Statistics are by one-way ANOVA with Tukey post-test (**B-F**) or by unpaired T-test (**G-H**), *p<0.05.

**Figure 8.**
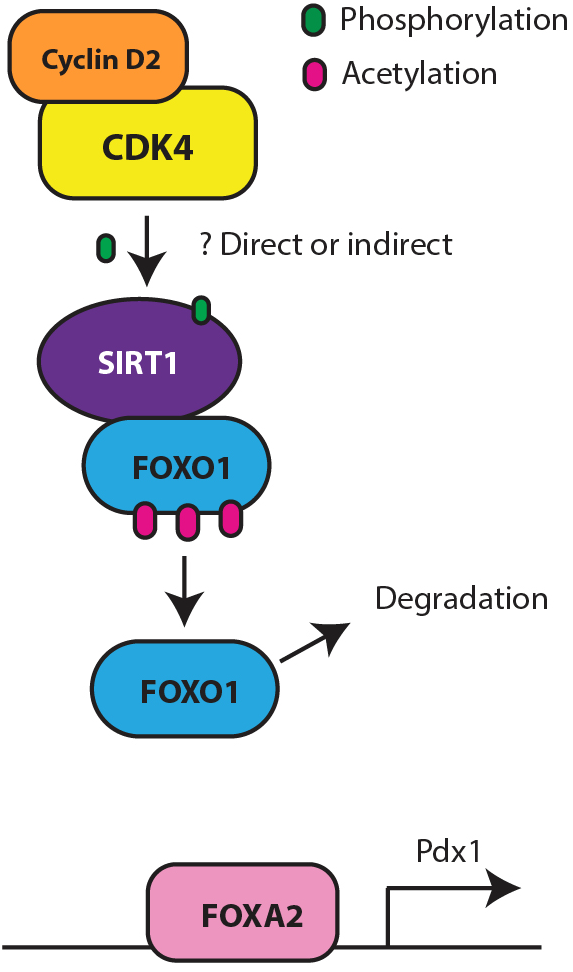
Model of how CDK4 may rescue FOXO1-mediated suppression of Pdx1 through deacetylation. Activation of the CDK4 kinase results in increased phosphorylation of the SIRT1 deacetylase, either through direct phosphorylation or indirectly, which deacetylates FOXO1, resulting in degradation of FOXO1 and derepression of *Pdx1*.

## DISCUSSION

This work identifies a new and surprising role for the cell cycle activator CDK4 to modulate insulin signaling in the adult pancreatic beta cell, counteracting FOXO1-mediated dedifferentiation. Replacing both endogenous alleles of *Cdk4* with *Cdk4-R24C* prevented diabetes in *Irs2* null mice by rescuing beta cell mass, function, and differentiation, restoring cytoplasmic FOXO1 location and nuclear PDX1 abundance in beta cells in vivo. Ex vivo, CDK4 inhibited FOXO1 suppression of *Pdx1* through two distinct mechanisms: AKT-dependent phosphorylation of FOXO1, and a second mechanism involving deacetylation, possibly by SIRT1. Together, these results demonstrate an unexpected role for CDK4 not only in driving the cell cycle but also in promoting beta cell differentiation and function.

The most important finding from these studies is that CDK4 can promote or maintain beta cell differentiation. It remains controversial whether beta cells must dedifferentiate to proliferate (10, 52). Replacing both alleles of *Cdk4* with *Cdk4-R24C* rescued not only beta cell proliferation but also dedifferentiation in diabetic *Irs2* null mice, supporting reports that it is possible to increase beta cell proliferation without negatively impacting beta cell function (52–57). A related observation was recently published in neurons (58). Since the primary source of new beta cells in the adult mouse is replication of mature beta cells (59, 60), the mutual incompatibility of proliferation and differentiation described in fetal development and adult stem cell activation paradigms may not apply. Indeed, CDKs are implicated in endocrine development (61, 62). Taken in this context, the observation that CDK4, a critical cell cycle driver in endocrine cells, can also promote their differentiation may represent a key piece in our understanding of adult islet homeostasis.

The degree of rescue of dedifferentiation by homozygous *CDK4-R24C* observed in this study was surprisingly complete, especially given the lack of improvement in insulin resistance, suggesting loss of CDK4/Cyclin D2 activity may be a primary cause of beta cell failure in IRS2 null mice. The IRS2 null phenotype was only partially rescued by FOXO1 haplosufficiency (13) or transgenic overexpression of *Pdx1* (27). *CDK4-R24C* may have additional parallel benefits such as bolstering beta cell mass through proliferation, or novel effects on lysosomal and mitochondrial biology (63). Our result is consistent with a prior observation that overexpression of CDK4-R24C under the insulin promoter rescued diabetes due to leptin receptor deficiency (64). That result was attributed to expansion of beta cell mass rather than a direct effect on beta cell differentiation. Other studies have also noted the surprising maintenance of physiological function of beta cells driven to proliferate by CDK4 activation (24).

These results reconfirm the known role for CDK4 in promoting beta cell proliferation (20, 21, 24). Insulin resistance in *Irs2-/-;Cdk4 R24C/R24C* mice likely contributed to increased proliferation (65, 66). Cyclin D2 drives beta cell compensation in response to insulin resistance (19), but islet Cyclin D2 levels are reduced in *Irs2*-null mice (14); rescue by *Cdk4-R24C* makes sense in this context. Residue 24 in CDK4 not only mediates INK family inhibition but also D-cyclin binding (67), so CDK4-R24C may have reduced regulatory inputs in both directions.

Proliferation was not statistically increased in *Irs2+/+;Cdk4 R24C/R24C* mice; increased beta cell mass in this group could be due to increased proliferation at an earlier timepoint. Alternatively, *Cdk4-R24C* allele may have expanded the pool of Ngn3+ endocrine precursors to increase beta cell mass in the adult (61, 62). Oddly, although proliferation was highest in *Irs2-/-; Cdk4 R24C/R24C* mice, beta cell mass was not higher in this group relative to *Irs2+/+;Cdk4 R24C/R24C* mice. Since *Irs2-/-;Cdk4 R24C/R24C* mice are insulin resistant with elevated fat mass, that group might have higher beta cell apoptosis, which was not assessed in this study, or competing suppression of proliferation by elevated free fatty acids (16). Finally, given the hyperinsulinemia in response to insulin resistance in *Irs2-/-;Cdk4 R24C/R24C*, CDK4 may synergize with activation of the unfolded protein response to drive beta cell proliferation (68).

Chronic hyperglycemia in *Irs2* null mice may contribute to dedifferentiation via oxidative stress or glucotoxicity, but it is likely not the only cause since *Pdx1* expression is reduced prior to the onset of hyperglycemia (27). Additionally, CDK4 promoted FOXO1 phosphorylation and increased *Pdx1* ex vivo in islet cells when glucose levels were held constant (Figures 4, 6 and 7). Surprisingly, *MafA* and *NeuroD1* were not induced by FOXO1 as previously reported (36), which suggests that FOXO1 targets may be sensitive to cellular context or type of stress (69).

These data suggest a role for INK family cell cycle inhibitors in beta cell dedifferentiation. Although the *Cdk4-R24C* allele has been termed ‘constitutively active’, the behavior of this mutant in our study was more consistent with a loss-of-function mechanism since the heterozygous state was indistinguishable from normal, and rescue was observed only in the homozygous state. Similarly, Rb phosphorylation was not always increased by *Cdk4*-*R24C* in the heterozygous state (70). A possible explanation is that this phenotype is dependent on loss of INK-family inhibition of CDK4, rather than activation of CDK4 kinase activity. Interestingly, p27, a Cip/Kip cell cycle inhibitor, is a cause of beta cell failure in *Irs2* null mice (71). The *R24C* mutation prevents INK4-family CDK4 inhibition (22, 23); in cancers, *Cdk4-R24C* mutation is synergistic with loss of function of Cip/Kip inhibitors (70) suggesting Cdk4-R24C may counteract a damaging combination of loss of Cyclin D2 with gain of p27 in IRS2 null beta cells.

Surprisingly, *CDK4-R24C* did not rescue insulin resistance in other metabolic tissues. CDK4 improves insulin sensitivity through PPARγ activation in adipocytes (25), suppression of hepatic glucose production (42) and by maintaining insulin signaling in adipocytes (26). Fat mass was increased in *Irs2-/-; Cdk4-R24C* mice, although this could have be secondary to hyperinsulinemia. In our study, fat mass was not increased in *Irs2+/+; Cdk4-R24C* mice. Since CDK4 acted through IRS2 in adipocytes (26) it is perhaps not surprising that Cdk4-R24C did not rescue insulin resistance in the context of IRS2 deletion.

Our data support a model in which FOXO1 subcellular localization is not the sole input into regulation of its gene targets. We initially expected that conditions in which FOXO1 immunostaining showed cytoplasmic localization would not show suppression of *Pdx1* mRNA, while nuclear FOXO1 would correlate with repressed *Pdx1* and other related changes in known FOXO1 targets (13, 29, 31, 36, 38). Our data at times did not support this model. Inhibiting nuclear export using leptomycin showed that even in glucose-excess conditions, FOXO1 passes through the nucleus. Despite forced retention of FOXO1 in the nucleus using leptomycin or AKT inhibition, *Pdx1* mRNA was not suppressed. Together, these experiments suggest that predominant nuclear localization of FOXO1 is not sufficient to suppress *Pdx1*, and that regulation of subcellular localization and transcriptional activity of FOXO1 is not as straightforward as the phosphorylated form being cytoplasmic and “off”, while the nuclear form is “on” and repressing *Pdx1*.

CDK4 increased phosphoryated FOXO1, but our data suggest CDK4 does not directly phosphorylate FOXO1 in islet cells. Besides AKT, other kinases such as SGKs, DYRK1 and CDK2 can phosphorylate FOXO1 (69, 72–74). CDK4 rescue of *Pdx1* gene expression was lost in the presence of AKT inhibitor, suggesting a requirement for AKT itself (most likely) or an AKT-dependent kinase. However, CDK4 rescued *Pdx1* expression even in the context of the unphosphorylatable FOXO1-ADA mutant (**Fig 6**), suggesting an additional mechanism besides phosphorylation.

CDK4 may regulate FOXO1 activity through deacetylation, since HAT inhibitors potentiated and HDAC inhibitors suppressed the rescue (36, 47–49). The data are consistent with CDK4 reducing acetylation of transcription factors rather than histones, since acetylation of histones tends to promote gene expression due to increased access of transcription factors to DNA (75). In line with this, CDK4 increased the phosphorylation and activation of SIRT1; sirtuins are the main class of HDACs that deacetylate FOXO1 (43, 44, 47, 49–51). However, we cannot conclude that the effects are via modulation of FOXO1 acetylation specifically, as we were not able to quantify acetylated FOXO1. FOXA2 promotes *Pdx1* expression (76) and can be deacetylated by SIRT1. FOXA2 deacetylation was reported to reduce its transcription activity in the liver and target it for degradation (77), but studies in the MIN6 beta cell line found that SIRT1 deacetylated FOXA2 to promote *Pdx1* expression (78). It will be interesting to investigate whether FOXO1 or FOXA2 acetylation is impacted by CDK4 in future studies.

Cyclin D2/CDK4 reduced FOXO1 protein abundance in cultured islet cells, consistent with accelerated degradation. While FOXO1 can be degraded after phosphorylation and cytoplasmic retention by 14-3-3 proteins (32, 79, 80), deacetylated FOXO1 is also more rapidly targeted for proteasomal degradation (36). Based on the observation that CDK4 promoted SIRT1 phosphorylation, we propose that CDK4/Cyclin D2 increases the deacetylation of FOXO1 via SIRT1, which leads to its degradation and derepression of *Pdx1*. In the path towards beta cell dedifferentiation, FOXO1 is nuclear during a “metabolic inflexibility” period (9, 81), but after exposure to chronic hyperglycemia, FOXO1 abundance declines (9). Since glucose increases Cyclin D2 protein expression (1, 14, 17), we speculate that the degradation of FOXO1 observed in late-stage dedifferentiation is mediated by chronic glucose stimulation of these cell cycle regulators and increased HDAC activity (36).

Strengths of this study include the use of parallel *in vivo* and *ex vivo* model systems to explore the fundamental biology of how a cell cycle regulator rescues diabetes phenotype in IRS2-null mice, a careful and comprehensive determination of the CDK4-mediated effect on FOXO1 biology and *Pdx1* expression, and the use of primary cells rather than transformed cell lines. Remaining questions include residual uncertainty as to which specific targets are phosphorylated by CDK4 to modulate the insulin signaling pathway, whether acetylation of FOXO1 itself is modulated, and whether these observations translate to human islets. Additionally, given the increased usage of CDK4/6 inhibitors in breast cancer, an important question is whether CDK4 inhibition impairs beta cell insulin signaling or differentiation.

In sum, we report the novel discovery that CDK4 promotes insulin signaling and FOXO1 degradation in the pancreatic beta cell, derepressing *Pdx1* expression and rescuing diabetes in IRS2-null mice. Our data suggest that CDK4 promotes both beta cell proliferation and differentiation, suggesting that if safe therapeutic approaches can be developed to increase beta cell mass via CDK4 they may preserve function.

## RESEARCH DESIGN AND METHODS

### Animal husbandry

*Cdk4-R24C* mice, in which the *R24C* mutation is knocked in at the *Cdk4* locus (21), were initially bred to *Irs2* heterozygous mice (B6;129-Irs2^tm1Mfw^/J) (12). Subsequently, mice heterozygous for both *Irs2* and *Cdk4-R24C* were bred and maintained on the C57BL/6N background. Mice were housed on a 12 hour light/dark cycle with ad libitum access to chow and water and studied at 8-12 weeks of age. Occasional malocclusion was present in this colony; non-diabetic mice with adult body weight of less than 20 grams were excluded from analysis. High fat diet pellets fed to female mice were 60% lard, obtained from Harlan/Envigo (TD.06414).

### Metabolic testing

Non-fasting blood glucose (ReliOn meter) and plasma insulin (Crystal Chem) were measured from tail tip blood samples. Intraperitoneal Glucose tolerance (GTT) and insulin tolerance (ITT) testing were performed as previously described (3). Mice were fasted for 5 hours prior to GTT (2g/kg; Hospira), while ITT (1.5U/kg; Humulin-R, Eli Lilly) was performed in the fed state. Body composition (fat/lean mass) was measured by the UMMS MMPC using 1H-MRS (Echo Medical System).

### Hyperinsulinemic-euglycemic clamps

Clamp studies were performed in the UMMS Mouse Metabolic Phenotyping Center (MMPC). Briefly, jugular vein catheters were implanted and mice were allowed to fully recover. Mice were fasted overnight, then primed with an initial insulin bolus (150mU/kg) followed by a continuous 2 hour infusion at a rate of 15pmol/kg/min. 20% glucose was infused at variable rates to maintain basal glucose levels, and blood glucose samples were taken at 10-20 minute intervals.

### Histology and Immunofluorescence

Mice were injected with BrdU (50g/kg i.p.) at 4 hours and 2 hours prior to euthanasia. Pancreata were dissected, fixed in 10% formalin (Sigma) for 4 hours, embedded in paraffin and cut in 5μm sections. Sections were stained as described (68); the antibodies used and unmasking conditions are detailed in **Suppl. Table 1**. Beta cell mass was quantified using full-pancreas sections histochemically stained for insulin (DAB) and hematoxylin and scanned. Mass was calculated as the product of the wet weight of the pancreas and the %insulin-labeled area quantified using Adobe Photoshop and Image J (16). Beta cell proliferation was expressed as the % insulin-labeled cells that colabeled for BrdU or pHH3 (16), using Cell Profiler pipelines developed and implemented by a blinded, unbiased lab member (82).

### Bulk RNA sequencing

Islets were isolated from male mice by ductal collagenase (Roche) injection and Ficoll (Histopaque-1077; Sigma) gradient as previously described (1). RNA was isolated using the Norgen All-in-one kit and sent to Quick Biology for library preparation and sequencing sequencing on the Illumina HiSeq 4000 with paired-end 150 bp reads. Paired-end reads were aligned to the mouse genome mm10 using the STAR aligner (83). Raw gene counts for each sample were generated from the subsequent bam files using HTSeq (84). Downstream analysis including filtering, normalization, and differential expression of the RNAseq counts was performed in edgeR (85). Principal component analysis and K-means clustering (2000 most variable genes, 10 clusters) were performed using the iDEP platform (86).

### Islet Isolation and Culturing

Islets were isolated as above from adult (10-40 week) C57BL/6J or C57BL/6N male or female mice and plated overnight in islet complete media (ICM: RPMI, 10% FBS (Sigma), penicillin/streptomycin, 5mM glucose). The following day islets were handpicked, trypsinized to single cells (0.05% trypsin), and plated on glass coverslips for immunofluorescence or plastic 24 well plates for protein/RNA analyses. One day after trypsinization, islet cells were exposed to adenovirus and/or drug for 72 hours in 15mM glucose unless otherwise specified. Adenoviruses were purchased from Vector Biolabs, and a list of all adenoviruses and multiplicity of infection (MOI) used is detailed **in Suppl. Table 2**. The FOXO1-ADA adenovirus was created from a plasmid (Addgene #12143) (87). Chemicals used include Leptomycin B (100nM; Sigma), MK-2206 (5μM; Selleckchem), C646 (25μM; Selleckchem), MB-3 (10μM; Santa Cruz Biotechnology), salermide (25μM; Tocris), vorinostat/SAHA (5μM; Selleckchem) and sodium butyrate (0.5mM; Selleckchem). For starvation experiments, islet cultures were cultured in 2mM glucose without FBS for 16 hours before harvesting. After the experiments, cells were fixed for immunofluorescence as described below, harvested for RNA and protein with SKP buffer with beta-mercaptoethanol per protocol (Norgen), or for protein with lysis buffer with PhosSTOP (Millipore-Sigma) as described (14).

### Quantitative PCR

RNA was isolated from dispersed islet experiments using the Norgen All-in-One kit. cDNA (SuperScript IV VILO; Thermo Fisher) was amplified (PerfeCTa SYBR; VWR) using either Eppendorf or BioRad thermocyclers (14) and analyzed by the ΔΔCT method. Primers are in **Suppl. Table 3**.

### Western Blotting

Immunoblots were performed as described (14). Briefly, dispersed islet cells were harvested either with the Norgen All-in-One kit or with Lysis Buffer (0.5M Tris pH 6.8, 10% SDS, 100mM DDT, 10mg/ml APMSF, protease inhibitor), both supplemented with PhosSTOP (Millipore-Sigma). Protein lysates were separated by SDS-PAGE, transferred to PVDF membrane (BioRad), and blocked in either 5% milk or BSA (w/v) in PBS with 0.1% Tween-20. Antibodies include pFOXO1 S256 (#84192, Cell Signaling Technology), FOXO1 (#2280, Cell Signaling Technology), pSIRT1 (JJ206-6; Novus) and Actin (MAB1501; Sigma). Data was collected on film or using a BioRAD gel doc station, using ECL (Thermo Fisher), ECL Prime (GE Healthcare), or SuperSignal West Femto (Thermo Fisher) and bands were quantified using ImageJ.

### Immunofluorescence

Dispersed mouse islets were fixed in 4% PFA for 10 minutes and stored in PBS at 4°C. Fixed cells on coverslips were immunostained as described (14). Briefly, coverslips were blocked in goat serum based block plus 0.1% Triton-X, then stained with the following antibodies: Insulin (1:200, Dako), and FOXO1 (1:50; Cell Signaling Technology), followed by Alexa Fluor secondary antisera (Invitrogen), and coverslips were mounted onto slides using Fluoroshield™ with Dapi (Sigma). Coverslips were imaged on either a Leica SP-5 Laser Scanning Confocal Fluorescence microscope, TissueGnostics Fluorescence Slide Scanner, or a Nikon TE2000-E2 inverted microscope. Quantification of nuclear FOXO1 was performed by a blinded individual utilizing a trinary scoring system.

### Statistics

Data were analyzed using GraphPad Prism. Most values are expressed as mean ±SEM except when specified otherwise. P values were calculated using *t* test, one-way ANOVA, or two-way ANOVA, as specified. P<0.05 was considered significant.

### Study Approval

All mouse studies were approved by the Institutional Animal Care and Use Committees of the Univerisity of Pittsburgh, University of Massachusetts Medical School, and Weill Cornell Medicine.

## Author Contributions

RES performed the majority of the experiments, contributed to study design and conceptualization, data analysis, data interpretation, funding acquisition, and manuscript preparation. RBS contributed to the *in vivo* metabolic studies and assisted with the RNAseq studies. HLG performed the western blots for the acetylation-related studies. CD assisted with image acquisition for the FOXO1 overexpression studies. DR performed the initial RNAseq data analyses. SR kindly provided the *CDK4-R24C* mice. LCA conceptualized the study and contributed to data analysis and interpretation, funding acquisition, and manuscript preparation. All authors had the opportunity to review and edit the manuscript.

## Acknowledgments

Financial support for this work was provided by NIDDK grants R01-DK124906 and R01-DK114686 to LCA; UMass MSTP 5T32GM107000 and NIDDK 5F30DK112591 to RES; the NIDDK Mouse Metabolic Phenotyping Centers (National MMPC, RRID:SCR_008997, www.mmpc.org) and specifically U2C-DK093000.

## FIGURE LEGENDS

**Supplemental Figure 1. Metabolic assessment of Irs2-null female mice without or with homozygous replacement of *Cdk4* with *Cdk4-R24C*. A**: Breeding dams and sires doubly heterozygous for *Cdk4-R24C* (whole body knock-in) and the *Irs2-*null allele (whole-body) produced littermate experimental mice of the genotypes shown. *Irs2 +/-* progeny were not studied. Panels **B-D** show data obtained from 14-week-old females fed with regular chow: 5-hour-fasting blood glucose (**B**) and blood glucose time course (**C**) or AUC (**D**) after intraperitoneal glucose challenge. Since *Irs2*-null females were not diabetic in normal chow conditions we performed similar experiments after 4 weeks of high fat feeding (**E-J**). 5-hour fasting blood glucose showed minimal hyperglycemia in *Irs2-/-* females on HFD (**E**). Glucose challenge identified hyperglycemia in *Irs2-/-; Cdk4-wt/R24C* females by 2-way ANOVA (**F**) but not by ANOVA of AUC (**G**). Body composition analysis of females after 4 weeks HFD by 1H-MRS Echo-MRI found no difference in body weight (**H**), % lean mass (**I**) or % fat mass (**J**). Number of replicates is shown for each panel except **C** (see **D**) and **F** (see **G**). Statistics are by one-way (**B, D, E, G, H, I, J**) or two-way (**C, F**) ANOVA with Tukey post-test. *p<0.05; **p<0.01; ***p<0.001; ****p<0.0001.

**Supplemental Figure 2. RNA sequencing of whole islets directly after isolation revealed numerous gene expression changes between groups. A**: Goes with the principal component analysis shown in Figure 3C. Intriguingly, PC1 is dominated by RNA processing related genes, and PC2 contains genes related to peptide biosynthesis. **B-D**: Scatterplots comparing *Irs2+/+; Cdk4-wt/wt* (Control) versus *Irs2-/-;Cdk4-wt/wt* (Diabetic) (**B**) or *Irs2-/-;Cdk4-wt/wt* (Diabetic) versus *Irs2-/-;Cdk4-R24C/R24C* (Rescue) (**C**) show numerous gene changes both upregulated and downregulated; changes are quantified in (**D**).

**Supplemental Figure 3. CDK4 overexpression increases FOXO1 phosphorylation through an indirect mechanism that requires AKT**. Mouse islet cells cultured in 15mM glucose were transduced with Ad-cre (control virus) or Ad-Cdk4 for 48 hours, followed by 24h exposure to the MK-2206 AKT inhibitor or vehicle. (**A**) Lysates were subjected to immunoblotting with antisera against phosphorylated FOXO1 (S256), phosphorylated AKT (S473), total AKT, or Actin. Quantification showed that CDK4 overexpression did not alter the ratio of p-AKT to total AKT (**B**), and that MK-2206 reduced p-AKT regardless of CDK4 overexpression (**B**). On the other hand, CDK4 overexpression markedly increased p-FOXO1 (**C**), but inhibition of AKT with MK-2206 completely prevented the CDK4-induced increase in p-FOXO1.

