## Supplemental figures for "Noncanonical CDK4 signaling rescues diabetes in a mouse model by promoting beta cell differentiation"

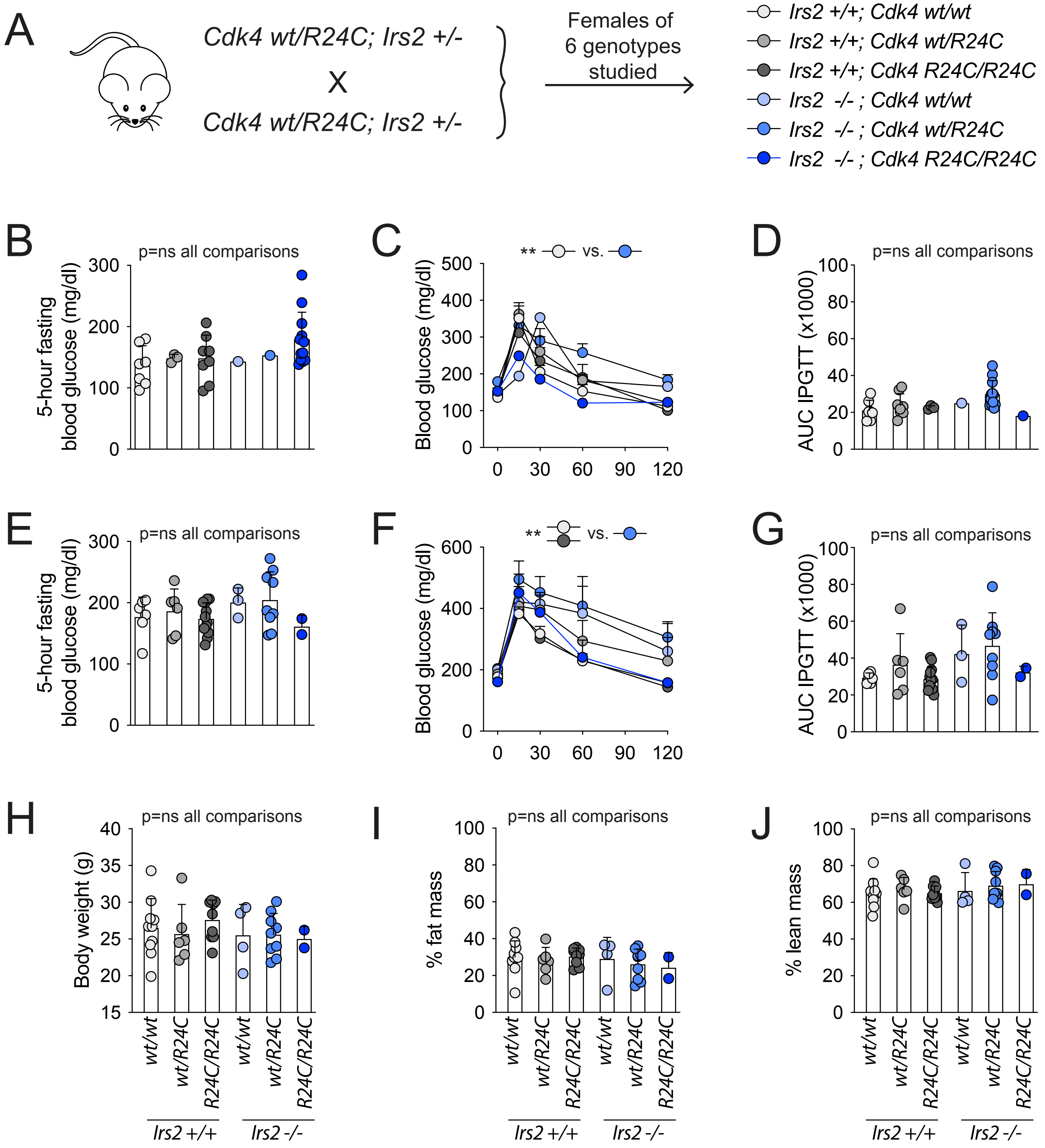


**Supplemental Figure 1. Metabolic assessment of *Irs2*-null female mice without or with replacement of *Cdk4* with *Cdk4-R24C*.** A: Breeding dams and sires doubly heterozygous for *Cdk4-R24C* (whole body knock-in) and the *Irs2*-null allele (whole-body) produced littermate experimental mice of the genotypes shown. *Irs2* +/- progeny were not studied. Panels B-D show data obtained from 14-week-old females fed with regular chow: 5-hour-fasting blood glucose (B) and blood glucose time course (C) or AUC (D) after intraperitoneal glucose challenge. Since *Irs2*-null females were not diabetic in normal chow conditions we performed similar experiments after 4 weeks of high fat feeding (E-J). 5-hour fasting blood glucose showed minimal hyperglycemia in *Irs2*-/- females on HFD (E). Glucose challenge identified hyperglycemia in Irs2-/-; *Cdk4-wt/R24C* females by 2-way ANOVA (F) but not by ANOVA of AUC (G). Body composition analysis of females after 4 weeks HFD by 1H-MRS Echo-MRI found no difference in body weight (H), % lean mass (I) or % fat mass (J). Number of replicates is shown for each panel except C (see D) and F (see G). Statistics are by one-way (B, D, E, G, H, I, J) or two-way (C, F) ANOVA with Tukey post-test. *p<0.05; **p<0.01; ***p<0.001; ****p<0.0001.

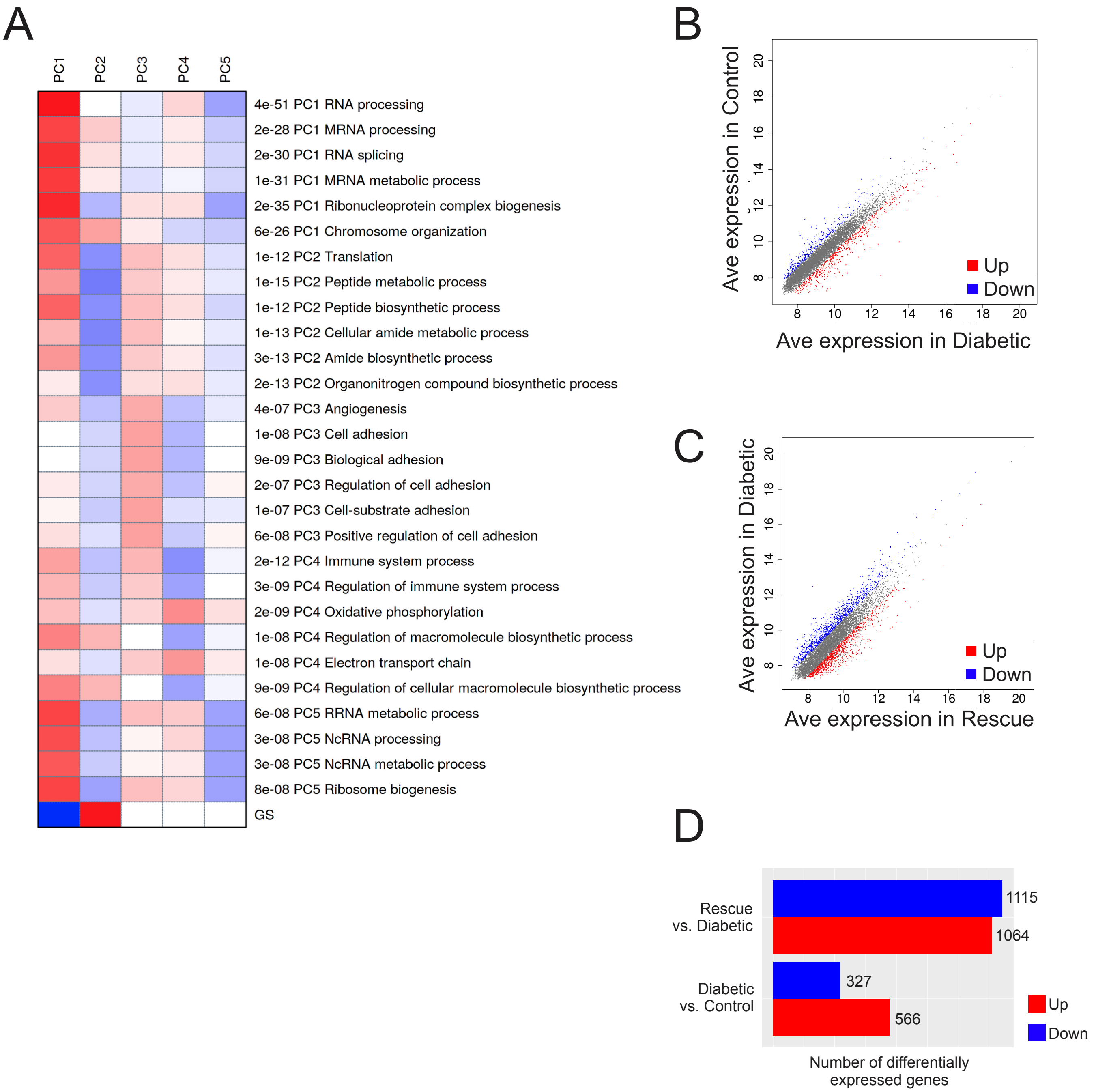


**Supplemental Figure 2. RNA sequencing of whole islets directly after isolation revealed numerous gene expression changes between groups.** A: Goes with the principal component analysis shown in Figure 3C. Intriguingly, PC1 is dominated by RNA processing related genes, and PC2 contains genes related to peptide biosynthesis. B-D: Scatterplots comparing Irs2+/+; *Cdk4-wt/wt* (Control) versus *Irs2-/-;Cdk4-wt/wt* (Diabetic) (B) or *Irs2-/-;Cdk4-wt/wt* (Diabetic) versus *Irs2-/-;Cdk4-R24C/R24C* (Rescue) (C) show numerous gene changes both upregulated and downregulated; changes are quantified in (D).

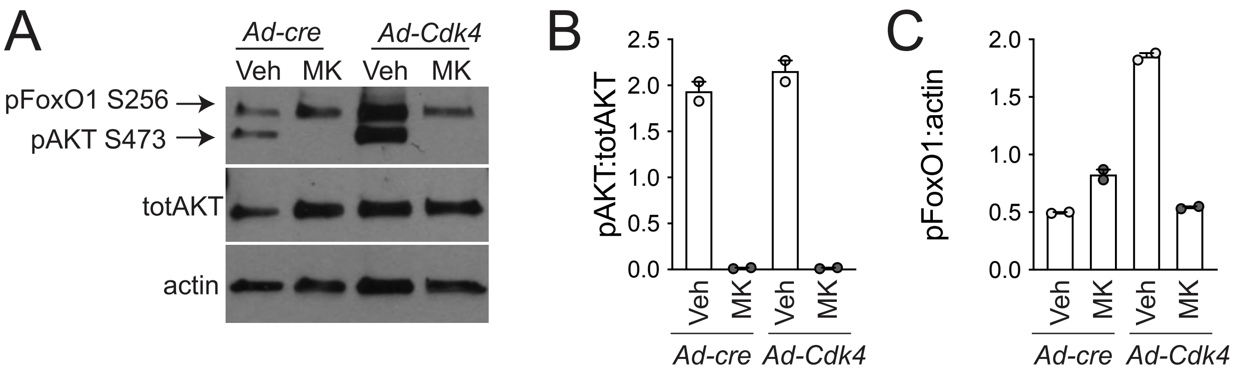


**Supplemental Figure 3. CDK4 overexpression increases FOXO1 phosphorylation through an indirect mechanism that requires AKT.** Mouse islet cells cultured in 15mM glucose were transduced with Ad-cre (control virus) or Ad-Cdk4 for 48 hours, followed by 24h exposure to the MK-2206 AKT inhibitor or vehicle. (A) Lysates were subjected to immunoblotting with antisera against phosphorylated FOXO1 (S256), phosphorylated AKT (S473), total AKT, or Actin. Quantification showed that CDK4 overexpression did not alter the ratio of p-AKT to total AKT (B), and that MK-2206 reduced p-AKT regardless of CDK4 overexpression (B). On the other hand, CDK4 overexpression markedly increased p-FOXO1 (C), but inhibition of AKT with MK-2206 completely prevented the CDK4-induced increase in p-FOXO1.
